# Southern Iberia as a hotspot of wild grapevine genetic diversity

**DOI:** 10.64898/2026.04.14.718376

**Authors:** Juan Jacobo Rodríguez-Felizzola, José Luis Blanco-Pastor, Jesús J. Soriano Bermúdez

## Abstract

**Aim:** The commercial interest of grapevines (*Vitis vinifera* L.) has prompted numerous studies on their origin and genetic resources in the context of global change. However, genomic-scale information on diversity patterns and genetic structure in southwestern Europe remains scarce. This study infers the genetic structure, gene flow events between genetic groups, and genetic refugia of *Vitis vinifera* ssp. *sylvestris* in the Iberian Peninsula.

**Location:** The Iberian Peninsula.

**Taxon:** The wild grapevine, *Vitis vinifera* L. ssp. *sylvestris*

**Methods:** We reanalyzed a set of 137 complete genomes of *V. vinifera* ssp. *sylvestris*. After variant calling, validation and annotation, we obtained a high-quality SNP dataset. Using these markers, we performed phylogenetic and population structure analyses to determine the number and spatial distribution of genetic groups and their contact zones. Next, we inferred the timing and directionality of gene flow events between groups. Finally, heterozygosity and allele rarity were estimated to identify populations with high conservation value.

**Results:** We detected three major ancestral populations and four putative genetic refugia in the south of the Iberian Peninsula. Demographic analyses indicate sustained gene flow between ∼21,000 and ∼7,000 years ago from a North African ancestral group into Iberian wild populations in the south. Heterozygosity and allele rarity analyses identified populations of high conservation value in a variety of areas within the Iberian Peninsula.

**Main Conclusions:** We identify the biogeographical factors behind the long-known singularity of wild Iberian grapevines. The southern Iberian Peninsula is a hotspot of genetic diversity for wild grapevines, hosting three ancestral populations and multiple contact zones that acted as micro-refugia. The current genetic variability of Iberian wild grapevines is best explained by natural, climate-driven gene flow between African lineages with Middle Eastern origin and Iberian groups. These contacts were favored by climatic conditions during the late Pleistocene (∼21,000 years) and early Holocene (∼8,300 years). Our results dismiss a significant anthropogenic influence during Neolithic domestication for explaining the genetic composition of Iberian wild grapevine genotypes.

## Introduction

The grapevine (*Vitis vinifera L*.) is recognized as a species of great sociocultural importance and is considered one of the most economically important fruit species globally (Vivier & Pretorius, 2002; OIV, 2025). The challenge facing viticulture today is to adapt this crop to a changing, more unstable, and drier climate, factors that are highly limiting at the southern edge of its distribution in Europe (Hannah et al., 2013; Sgubin et al., 2023; van Leeuwen et al., 2024). Additionally, domestication-driven selection for productivity traits has led to genetic erosion of stress-resistance mechanisms, increasing the susceptibility of cultivated grapevines to pathogen infections and environmental constraints (Dolferus, 2014; Marrano et al., 2018). The wild grapevine (*Vitis vinifera* L. ssp. *sylvestris*, hereinafter *V. sylvestris*) is a genetic reservoir that has not been sufficiently explored and utilized in modern viticulture. Wild grapevine populations possess a wealth of adaptive traits shaped by millennia of natural selection under fluctuating and often harsh environmental conditions (Daldoul et al., 2023), resulting in populations that harbor great genetic diversity and significant agronomic and oenological potential. These properties make wild grapevines highly valuable as a genetic resource with potential use in the improvement of modern cultivated varieties (FAO., 2014; Petitpierre et al., 2023; Daldoul et al., 2025). This diversity includes desirable traits related to fruit quality and tolerance to abiotic and biotic stress, which currently represent key breeding objectives (Grassi & Arroyo-Garcia, 2020; Li & Gschwend, 2023; Daldoul et al., 2025). Given the edaphic and climatic diversity of the Iberian Peninsula, it is to be expected that the genetic diversity present in the wild grapevine populations from Spain and Portugal is a rich source of variability (Ocete et al., 2014; Arroyo-Garcia et al., 2016) that can be used to improve resistance to disease and climatic stress, the productivity, or even the quality of the wine of modern cultivated grapevines (De Andrés et al., 2012; Daldoul et al., 2025). Understanding the evolutionary and demographic history of wild grapevines is therefore essential to effectively harness this diversity for breeding and conservation.

The evolutionary history of wild grapevines has received special attention since the beginning of the 21st century, from single markers to genome-wide phylogeographic studies (Myles et al., 2011; Liang et al., 2019; Dong et al., 2023). Plastid DNA analyses showed a differentiation between western and eastern wild grapevines and a chlorotype (A) characteristic of wild populations on the Iberian Peninsula was identified (Arroyo-García et al., 2006). Microsatellite markers (SSR) allowed the identification of two genetic clusters of wild grapevines in Spain, one population distributed in the north and another in the south of the country (De Andrés et al., 2012). SSRs also reported introgression from wild populations into cultivated varieties, but not in the opposite direction (Xiao et al., 2023). This asymmetry was also observed using 5,387 single-nucleotide polymorphisms (SNPs), which also identified the genetic uniqueness of wild grapevines in southern Spain, markedly different from other European populations (Myles et al., 2011). The latter resequencing of complete genomes of cultivated and wild grapevines has shown a clearer differentiation between western and eastern genotypes across their distribution range and found some genetic similarity between wild and cultivated populations in the Iberian Peninsula (Liang et al., 2019), thus supporting the introgression detected by SSR markers (De Andrés et al., 2012).

Recent analyses, also using complete genomes, have confirmed the differentiation between western and eastern varieties, as well as the introgression mentioned above (Freitas et al., 2021; Magris et al., 2021). In addition, Freitas et al., (2021) reported gene flow from wild populations to cultivated varieties again, which is particularly pronounced within the Iberian Peninsula, giving this region a unique genetic character among cultivated grapevines. Finally, Dong et al., (2023) conducted an exhaustive study of more than 3,000 complete genomes from across the Mediterranean basin, which allowed them to trace the history of large-scale domestication and the relationships between cultivated and wild grapevines. Here, they confirm the continuous geographical and genetic differentiation of wild grapevines from east to west, and the independent events of introgression between wild and cultivated varieties in different regions of the Mediterranean.

The Iberian Peninsula has been a genetic refugium for this and other wild species during the last glaciation (Levadoux, 1956; López de Heredia et al., 2007; Arroyo-García et al., 2016). During the Last Glacial Maximum (∼21,000 years ago), the ecotypes of eastern wild grapevines (East Asia and the Caucasus) and western wild grapevines (Central Europe and the Iberian Peninsula) completely differentiated their ecological niches because of geographical isolation (Dong et al., 2023). Phylogeographic studies of wild grapevines have been conducted on a continental scale (Zhou et al., 2017; Liang et al., 2019; Sivan et al., 2021; Dong et al., 2023). These analyses, undertaken on a large geographical scale, have not traced regional events on a smaller scale, particularly differentiation and introgression events within the Iberian Peninsula, where only regional studies based on SSRs or limited sampling have been performed so far (Arroyo-García et al., 2016; Loureiro et al., 2023). These studies have shown a north-south differentiation in wild grapevines (De Andrés et al., 2012; Dong et al., 2023), but the phylogeographic boundaries of these lineages, together with the splits and admixture events, remain unknown. Here, we infer the genetic structure and gene flow events between *V. sylvestris* individuals at a small regional scale (Iberian Peninsula), identify the geographical areas that have acted as genetic refugia for the species, and investigate whether anthropogenic activities have influenced its demographic history. Our working hypotheses are that: i) genomic analyses on the scale of the Iberian Peninsula will enable us to identify additional genetic groups beyond the north-south genetic differentiation; ii) specific contact zones and individuals with high genetic diversity and rarity will be identified for conservation purposes; and (iii) grapevine domestication was not key for shaping the wild grapevines genetic structure in the Iberian Peninsula.

## Material and Methods

### Data acquisition

The *V. sylvestris* genome sequences from Dong et al., (2023) were analyzed and are available in the online repository “China National Center for Bioinformation”. A total of 137 sequences (See Table S1 in Appendix S1) were downloaded, of which 117 are located in the Iberian Peninsula (92 in Spain and 25 in Portugal). For the genetic clusters defined by Dong et al., (2023) corresponding to *V. sylvestris* lineages K1 (Central Europe), K2 (Western Asia), K6 (Caucasus), and K8 (Iberian Peninsula), five representative sequences were selected from each, with a membership percentage of more than 80% for each of these groups.

### Variant calling, validation, and annotation

Sequences were filtered with FASTP (v 0.24.0) (Chen et al., 2018). Subsequently, using the VS-1 genome as a reference, the sequences were aligned with BWA-MEM2 (v.2.2.1) (Vasimuddin et al., 2019), with the default parameters. Samtools (v.1.21) (Danecek et al., 2021) and Picard (v.3.3.0; https://broadinstitute.github.io/picard) were used to sort the aligned sequences and remove duplicates, respectively. Additionally, the sequencing depth, duplication rate, and mapping percentage for each sequence were calculated using BAMDST (v.1.1.0; https://github.com/shiquan/bamdst). Values that did not meet the mean ± 3S.D. of the evaluated parameters were considered outliers and discarded for further analysis, following the methodology of Dong et al., (2023). The final dataset included 131 sequences of *V. sylvestris*. Finally, a sample of *V. riparia* (SJAQ01.1) (Girollet et al., 2019) was inserted as an outgroup.

The chromosomes of the *V. sylvestris* VS-1 genome (Dong et al., 2023) were used as a reference for the identification of single nucleotide polymorphisms (SNPs) and insertion/deletion polymorphisms (Indels), without taking into account unanchored sequences (scaffolds). GATK4 (v.4.6.1.0) (Van der Auwera & O’Connor, 2020) was used to obtain these variants, following the pipeline described by Van der Auwera et al., (2013). First, variant calling was performed for each of the sequences using GATK HaplotypeCaller; then, a joint genotyping analysis of the gVCF (genomic Variant Call Format) was carried out. For the filtering of SNPs and Indels, the parameters indicated by Dong et al., (2023) were replicated, these being: for SNP filtering: “QD<2.0, QUAL<30.0, SOR>3.0, FS>60.0, MQ<40.0, MQRankSum<-10.0, ReadPosRankSum<-8.0.” And for Indel filtering: “QD<2.0, QUAL<30.0, SOR>5.0, FS>100.0, InbreedingCoeff<-0.8.”

### Phylogenetic analyses and genetic structure

For phylogenetic and genetic structure analyses, the approach of Dong et al., (2023) was followed. To reduce the dataset to a set of data that could be used for phylogenetic analysis, PLINK (v1.90b6.24) (Chang et al., 2015) was used to filter the SNP dataset, eliminating those with more than 20% missing data and with a minor allele frequency (MAF) of less than 0.05. In addition, SNPs with high linkage disequilibrium (LD) (r2 >= 0.5) within a continuous window of 50 steps were removed. The resulting SNP dataset was processed with SNPhylo (v. 20180901) (Lee et al., 2014). The result obtained in phylip format was used to construct a maximum likelihood (ML) phylogenetic tree with RAxML NG (v1.2.2) (Kozlov et al., 2019) using 32 random search trees and 100 TBE bootstraps (Lemoine et al., 2018). The phylogenetic tree was visualized in FigTree 1.3.1 (Rambaut, 2010). We estimated genetic ancestry and the number of ancestral populations (K) with ADMIXTURE (v.1.3.0) (Alexander et al., 2009), we used a cross-validation (CV) test, which was run with its default parameter of 5 folds (Alexander & Lange, 2011).

To determine migration events based on allele frequencies, an analysis was performed in Treemix (v.1.13) (Pickrell & Pritchard, 2012). To simplify the analysis and following the methodology of Dong et al., (2023), individuals with at least 75% ancestry for each of the ancestral populations determined in the previous step were used. Additionally, migration events were also evaluated with individuals who had more than 70% ancestry. Both sets of SNPS were filtered for missing data and monomorphisms, since the analysis only accepts biallelic SNPS. The optimal number of migrations present between ancestral populations was determined using the R “OptM” package (v.0.1.6) (Fitak, 2021), evaluating between 1 and 10 migration events (m). To create the consensus tree, *V. riparia* was established as the outgroup, and Treemix was run with bootstrap 1000 -k 500. For each migration event, the tree was constructed 10 times using random seeds.

Finally, we analyzed the demographic history of ancestral populations with Momi2 (v.2.1.21) (Kamm et al., 2020). We evaluated ten demographic models (Table 1) that considered the results of Treemix. These models were evaluated to determine whether the current demographic history of wild grapevines in the Iberian Peninsula has been influenced by migration events determined by a climatic context in which wild grapevine populations came into contact during the Pleistocene in the Last Interglacial period (∼130,000 years ago) or the Last Glacial Maximum (∼21,000 years ago) (Dong et al., 2023). Or whether, on the contrary, the current genetic structure has been determined by migrations caused by anthropogenic activities related to the development of agricultural models and the domestication of grapevines by early Neolithic societies ∼7,000 years ago (Myles et al., 2011; De Andrés et al., 2012; McGovern et al., 2017; Magris et al., 2021; Dong et al., 2023). For this purpose, and following Dong et al., (2023), the five individuals with the highest ancestry from each genetic group were selected, and SNPs were filtered out for coding and repetitive regions. The spectrum of frequencies of folded sites, without knowledge of the ancestral allele (folded SFS), was divided into 100 blocks of equal size for jackknifing and bootstrapping. A single gene flow event and a constant population size were assumed for the four populations (K1-K3 and outgroup). For each of the models evaluated, the Akaike information criterion (AIC) was calculated, and the model with the lowest AIC value was selected.

**Table 1.** Demographic models generated with Momi2 to evaluate alternative migration scenarios among wild grapevine populations from the Iberian Peninsula and North Africa. K1 corresponds to the north of the Iberian Peninsula, K2 in the south of the Iberian Peninsula, and K3 in North Africa. The arrows indicate the direction of gene flow. The models include single migration pulses that occurred 7 ka, 21 ka (Last Glacial Maximum, LGM), and 130 ka (Last Interglacial, LIG), as well as scenarios of two pulses at 7 and 21 ka and multi-period models that include pulses at 7, 21, and 130 ka. These alternative scenarios were designed to test whether Pleistocene climate fluctuations facilitated episodic or sustained connectivity between populations in North Africa and the Iberian Peninsula. logL = Likelihood. K = number of parameters evaluated. AIC = Akaike information criterion. ΔAIC = difference between the AIC of a given model and the AIC of the best model.

| Model | Migration direction | Migration type | N. of pulses (time period considered) | logL | K | AIC | ΔAIC |
| --- | --- | --- | --- | --- | --- | --- | --- |
| 1 | K3 → K1 and K2 | Single pulse | 1 (~ 7 ka) | -45349900.21 | 3 | 90699806.41 | 10056.08 |
| 2 | K3 → K1 and K2 | Single pulse | 1 (~ 21 ka) | -45349904.37 | 3 | 90699814.75 | 10064.42 |
| 3 | K3 → K1 and K2 | Single pulse | 1 (~ 130 ka) | -45358151.62 | 3 | 90716309.24 | 26558.91 |
| 4 | K1 and K2 → K3 | Single pulse | 1 (~ 7 ka) | -45349901.64 | 3 | 90699809.28 | 10058.94 |
| 5 | K1 and K2 → K3 | Single pulse | 1 (~ 21 ka) | -45349904.03 | 3 | 90699814.06 | 10063.73 |
| 6 | K1 and K2 → K3 | Single pulse | 1 (~ 130 ka) | -45358157.06 | 3 | 90716320.12 | 26569.79 |
| 7 | K3 → K1 and K2 | Continuous migration | 2 (~ 7 - 21 ka) | -45344870.17 | 5 | 90689750.33 | 0 |
| 8 | K1 and K2 → K3 | Continuous migration | 2 (~ 7 - 21 ka) | -45349903.19 | 5 | 90699816.37 | 10066.04 |
| 9 | K3 → K1 and K2 | Continuous migration | 3 (~ 7 - 21 - 130 ka) | -45352623.22 | 7 | 90705260.44 | 15510.11 |
| 10 | K1 and K2 → K3 | Continuous migration | 3 (~ 7 - 21 - 130 ka) | -45358147.88 | 7 | 90716309.77 | 26559.43 |

### Analysis of heterozygosity and allelic rarity

The genetic rarity of each individual was estimated based on the allele frequencies of the SNPs obtained with PLINK (v1.90b6.24) (Chang et al., 2015). From these, a weight file was generated in which each allele was assigned a value inversely proportional to its frequency, defined as the negative logarithm to base 10 of the minor allele frequency (MAF), so that rarer alleles received a higher weight. Subsequently, the weights of the alleles present in each individual were added together, thus obtaining a cumulative index of individual genetic rarity. This value reflects the proportion of rare variants carried by each individual, allowing for comparison of the representation of rare alleles within the sample set. Heterozygosity per individual was also calculated using VCFtools (0.1.17) (Danecek et al., 2011).

## Results

After calling and filtering variants in the analyzed sequences, a dataset of 7,969,329 SNPs and 5,031,438 Indels was obtained. The Maximum Likelihood (ML) analysis (Figure 1) showed the formation of three main clades (see below), of which the clade consisting mainly of individuals predominantly from the blue group (K3, North Africa) that occupies a basal position relative to the other two clades being more derived (K2 and K1, south and north of the Iberian Peninsula). However, because the statistical support for the ML phylogenetic tree in its internal branches is low, the interpretation of the evolution of *V. sylvestris* based on this phylogeny is limited.

**Figure 1.**
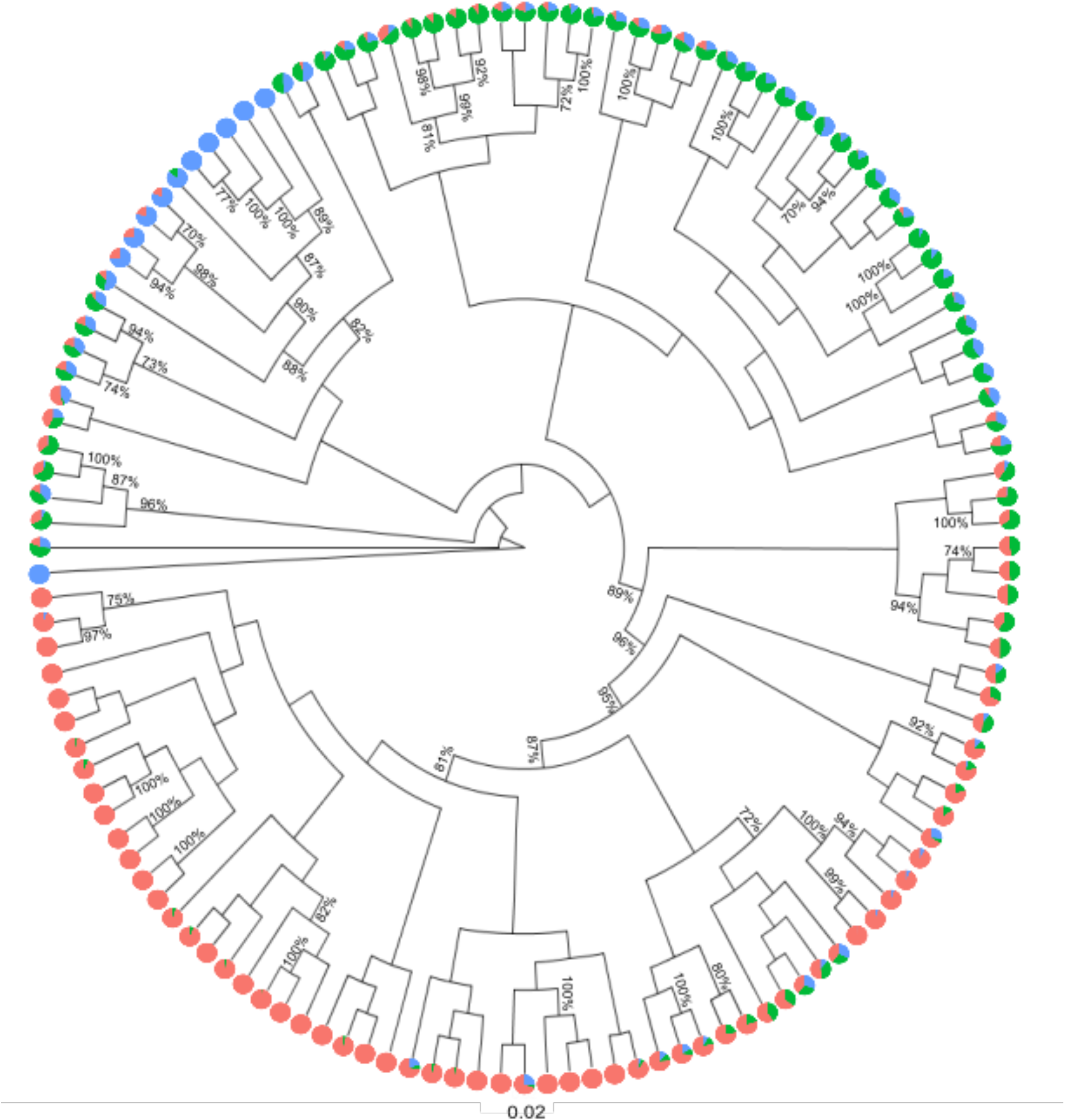
Maximum likelihood (ML) phylogeny of *V. sylvestris*. Values above 70% ML are indicated. The genetic ancestry of each individual is represented by circles at the tips of each branch. Red refers to K1 (northern Iberian Peninsula), green to K2 (southern Iberian Peninsula), and K3 (northern Africa) is represented by blue.

The CV test determined that K=3 was the optimal number of genetic groups, reaching its minimum absolute error at this value (See Figure S1 in Appendix S2). The admixture analysis for K=3 (Figure 2A) shows a higher proportion of individuals with K1 ancestry (hereinafter, northern Iberian Peninsula, NIP), followed by K2 (southern Iberian Peninsula, SIP) and finally K3 (North Africa, NA). With 64, 38, and 14 sequences, respectively. Colors can be related to the geographical location of the individuals (Figures. 2B and 2C). In the case of the NIP ancestral population (red), its dominance in the north is evident. In the south, on the other hand, no ancestral population is observed to dominate overwhelmingly, with decreasing NIP ancestries towards the south and intermediate individuals with ancestry from both the SIP and NA populations in the southern extreme of the peninsula. Meanwhile, for the sequences located in Portugal and south-central Spain, all three genetic ancestries are present and share dominance. The NIP-SIP combination is more notable in Portugal.

**Figure 2.**
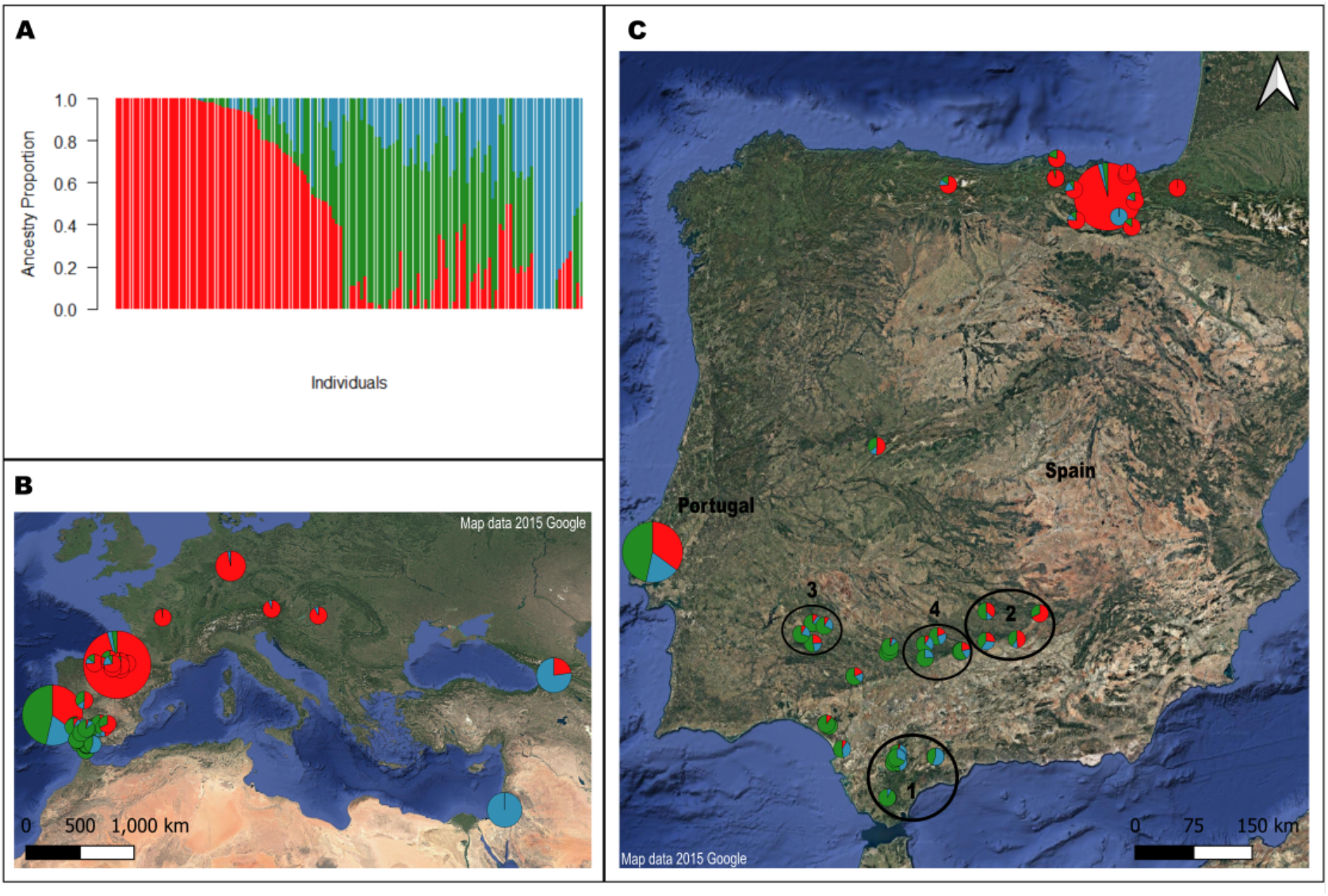
Population structure and geographic distribution of the *V. sylvestris* individuals analyzed. (A) Admixture. Red refers to K1 (north of the Iberian Peninsula), green to K2 (south of the Iberian Peninsula), and K3 (north of Africa) is represented by blue. (B and C) Geographic representation of the individuals analyzed. Genetic ancestry is included. The size of the circles is proportional to the number of individuals per location, from n=28 to n=1 (precise location of Portuguese and Basque Country accessions were unknown). Red refers to K1 (north of the Iberian Peninsula), green to K2 (south of the Iberian Peninsula, SIP), and K3 (north of Africa, NA) is represented by blue. The geographical areas of interest are included: 1) Los Alcornocales, 2) Sierra de Andújar, 3) Sierra de Aracena, and 4) Sierra deHornachuelos. Map projection: Mollweide equal area (EPSG:54009), equatorial scale.

Within the Iberian Peninsula, it is possible to identify the geographical areas where ancestral populations have come into contact, as evidenced by the change in the major ancestry of individuals. We can identify the areas of contact between two or three genetic groups and the precise delimitation of the contact zone (Figure 2C). In (1) Los Alcornocales, individuals with major K2 ancestry (green) are observed to the west of the mountain range and individuals with major K3 ancestry (blue) to the east of the mountain range, with no presence of K1 ancestry (red) delineating a clear K2–K3 boundary. In (2) Sierra de Andújar, individuals are mainly K2 (green) west of the mountain range and K1 (red) east of the mountain range, with a lower presence of K3 (blue). The Sierra de Aracena (3) and Sierra de Hornachuelos (4) exhibit high representation of the three genetic ancestries, emerging as hotspots of tripartite admixture.

In the north of the Iberian Peninsula, NIP ancestry is very dominant; however, it is important to highlight the presence of one individual who is entirely K3 (blue, SIP). We believe that this individual (CRR498555) could be a later introduction (human-mediated migration or bird dispersal) or a cataloging error in the database, given the difference with nearby individuals, warrants further verification of sampling coordinates and passport data (Figure 2C).

For the Treemix analysis, the optimal number of migration events (m) was m=1, which is evidenced by a notable increase in the average likelihood and explained variance between m=0 and m=1 (Figure S2 in Appendix S2). A phylogenetic tree for individuals with ≥ 75% ancestry from ancestral populations (Figure 3A) shows the phylogenetic tree resulting from Treemix, with a migration edge from NA (K3) to SIP (K2), indicating a gene flow event across the strait of Gibraltar, with a migration weight of 0.123569. This means that approximately 12% of the genetic pool of SIP (K2) comes from NA (K3). Similarly, this analysis allowed us to identify that the population NIP (K1) derives from the population SIP (K2), suggesting a south-north migration of the wild grapevine populations present in the Iberian Peninsula. However, A phylogenetic tree for individuals with ≥ 70% (Figure 3B) presents an opposite scenario, in which gene flow would be represented by a migration edge from an ancestral lineage NIP (K1) and towards NA (K3), with a migration weight of 0.0723, thus indicating that 7% of the genetic reservoir of NA (K3) is derived from NIP (K1). This would indicate a genetic migration of wild grapevine in a north-south direction. This scenario is less consistent with the geographic location of the ancestral groups, given that this contact must bypass K2, located between K1 and K3. In both cases, based on the drift parameter, it can be observed that the NIP population (K1) has accumulated greater genetic drift than the other ancestral populations evaluated.

**Figure 3.**
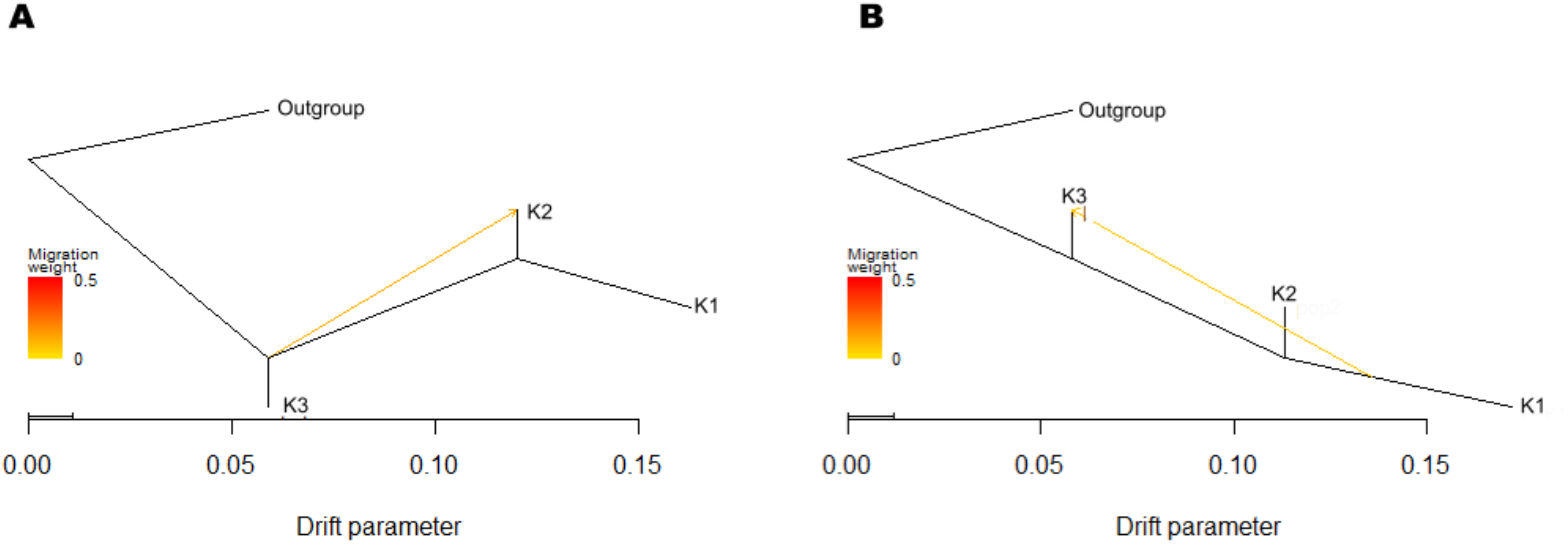
(A) Treemix phylogenetic tree for *V. sylvestris* individuals with ≥ 75% ancestry from ancestral populations. (B) Treemix phylogenetic tree for *V. sylvestris* individuals with ≥ 70% ancestry from ancestral populations. The migration weight gradient is included. The X-axis shows the genetic drift parameter. Population K3 refers to north of Africa (NA), K2 to south of the Iberian Peninsula, (SIP), and K1 to north of the Iberian Peninsula (NIP).

The results of Momi2 (Table 1) indicate that, with a logL of −45344870.17, model 7 is the one that best fits the data and explains the directionality of migration between the ancestral populations evaluated here. This model proposes that the ancestral NA (K3) wild grapevine population migrated to the NIP (K1) and SIP (K2) populations (Figure 4). Similarly, the model suggests that migration likely occurred continuously between ∼21,000 years ago and ∼7,000 years ago, rather than in a specific time pulse. Likewise, Figure 4 shows that the migration pulse is stronger (10.5%) at the beginning of the gene flow from K3 to K1/K2 than in recent times, where the migration pulse is 0.9%. The Momi2 results would support the model and direction of gene flow obtained in Treemix using individuals with ≥ 75% ancestry from ancestral populations (Figure 3A), where the population originating in Africa (NA) contributes with gene flow to the Iberian populations (NIP and SIP).

**Figure 4.**
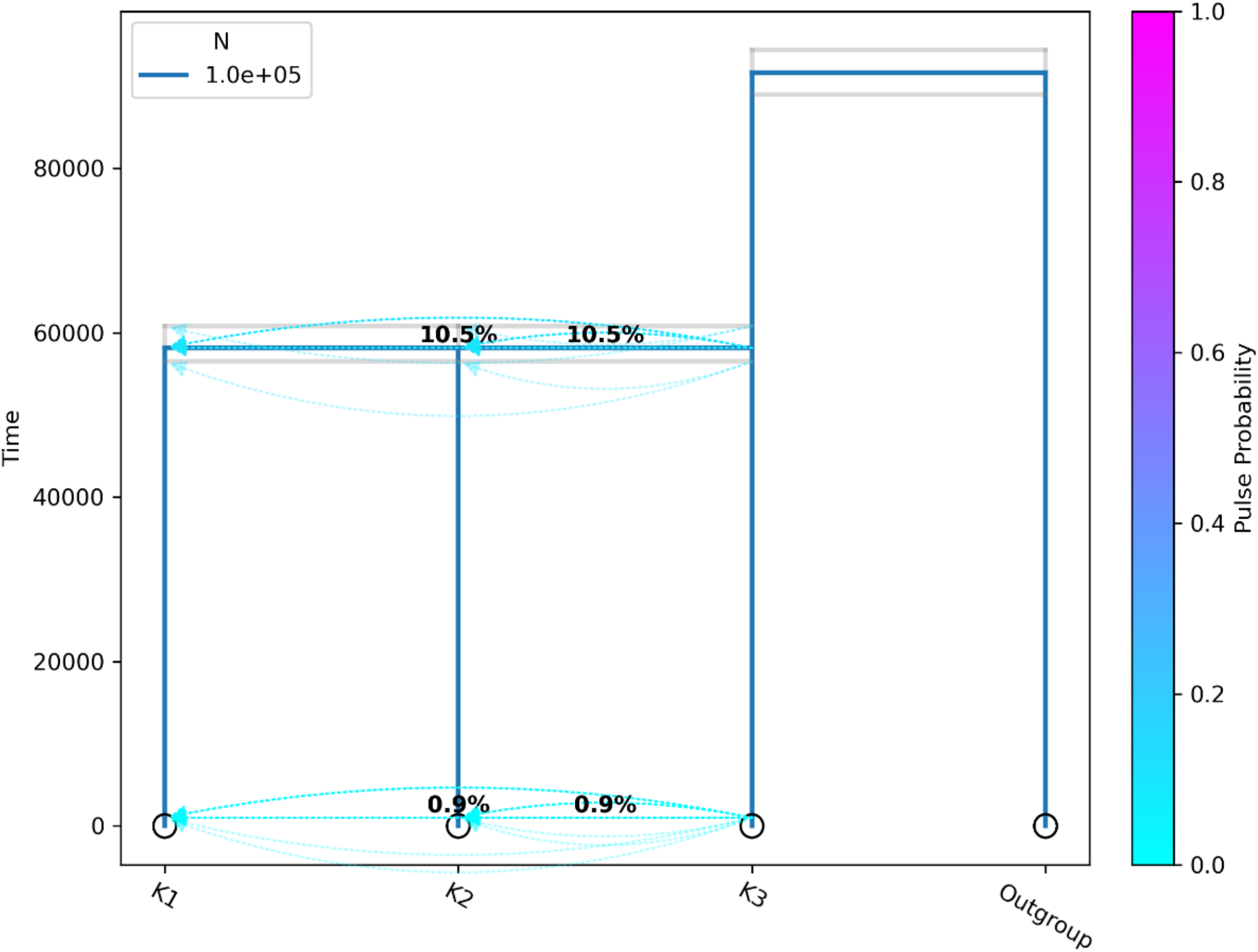
Momi demographic model. The arrows indicate the direction of migration of the *V. sylvestris* populations, and their color indicates the probability of the pulse; 10.5% 58,230 (55,460 ± 61,984) years ago and 0.9% 1,000 (1,000,45321 ± 1,000,45328) years ago. Population K3 refers to north of Africa (NA), K2 to south of the Iberian Peninsula, (SIP), and K1 to north of the Iberian Peninsula (NIP).

Finally, heterozygosity and allele rarity analyses indicate that individuals CRR498688, CRR499573, CRR498475, CRR498686, and CRR498859, in this order (Figure 5), are the individuals that combine the least frequent alleles and the highest heterozygosity. These individuals were not located in a single geographical area but spread across the Iberian Peninsula in multiple sampling sites.

**Figure 5.**
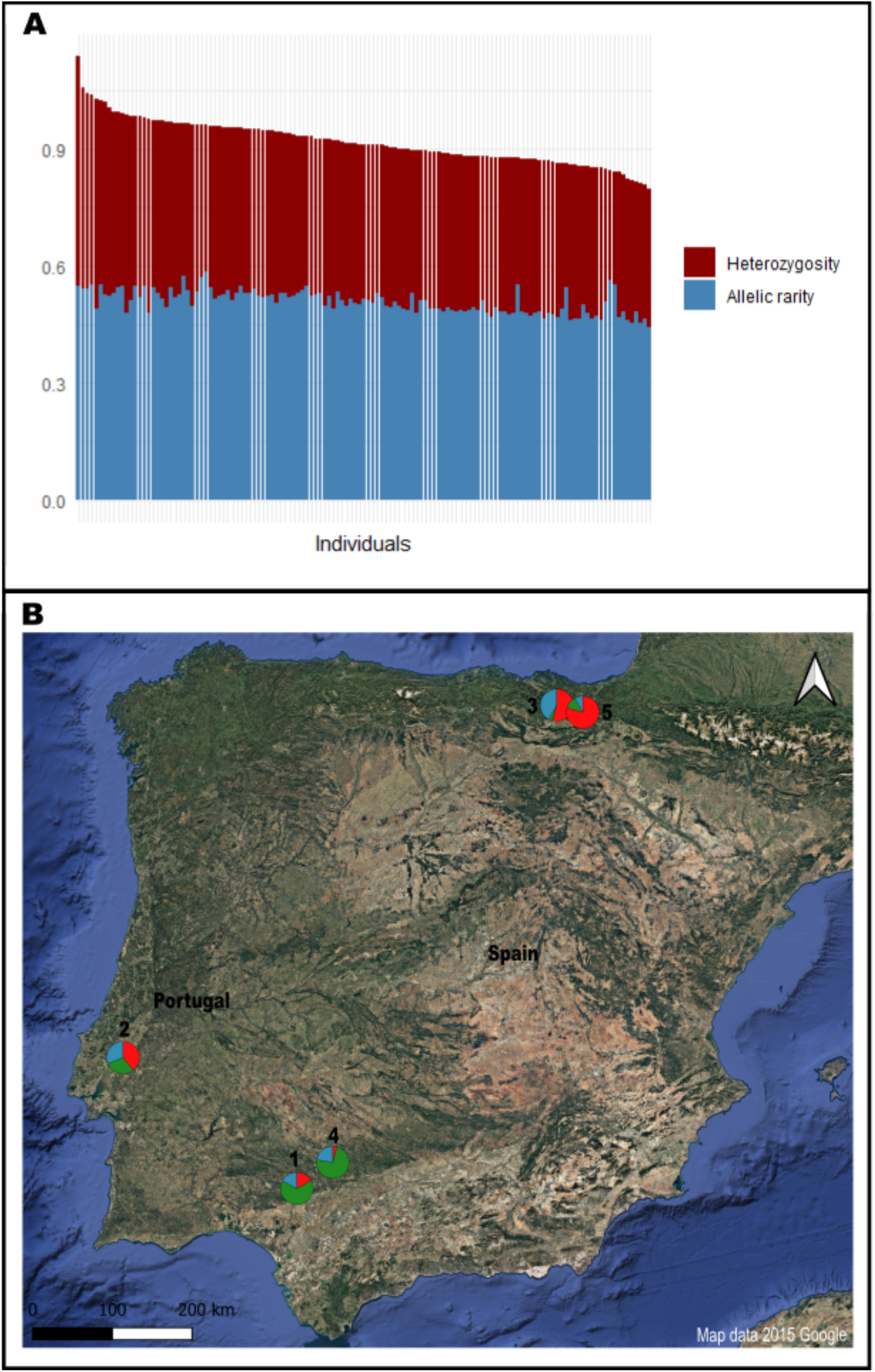
Heterozygosity and allele rarity analyses. (A) Total sum of allelic rarity (blue) and heterozygosity (red) values per individual. (B) Geographic representation of individuals with the least frequent alleles and the highest heterozygosity; 1) CRR498688, 2) CRR499573, 3) CRR498475, 4) CRR498686, and 5) CRR498859. Red refers to K1 (north of the Iberian Peninsula, NPI), green to K2 (south of the Iberian Peninsula, SIP), and K3 (north of Africa, NA) is represented by blue. Map projection: Mollweide equal area (EPSG:54009), equatorial scale.

## Discussion

The evolutionary history of *V. vinifera* in terms of its differentiation and domestication processes has been extensively studied at the continental and global levels (Grassi & De Lorenzis, 2021; Dong et al., 2023). However, to date, the study of the genetic structure and demography of *V. sylvestris* within the Iberian Peninsula has been limited. In the present study, the presence of three clusters or ancestral populations (K=3) within the region was identified, which expands on the work by De Andrés et al., (2012) and Freitas et al., (2021), who had reported two clusters (K=2) of Iberian origin, associated with a north-south geographical differentiation. The results of this study confirm the presence of this spatial pattern in the Iberian Peninsula but also reveal an east-west differentiation. Furthermore, rather than two, the presence of three ancestral populations is demonstrated, one of which is of African origin.

Further research is needed into the relationship between genotypes from the Middle East and North Africa, and to increase the sampling effort in the central Iberian Peninsula (central Spain) in order to clarify the patterns of gene flow across ancestral populations (Cantos et al., 2017). It is also essential to refine and obtain precise information regarding the collection localities of the Portuguese and the Basque Country samples. Together, these efforts would improve our understanding of the number of genetic groups, their geographical limits, and their relationships, help to clarify gene flow dynamics, and enhance the resolution of refugial areas and contact zones among the ancestral wild grapevine populations in the region.

Despite these limitations, we identified the greatest genetic diversity in the southern part of the Iberian Peninsula, with representative percentages of the three ancestral populations identified for the study area. In contrast, individuals from the north of the Iberian Peninsula are more homogeneous, as the dominance of NIP cluster (K1) is very high, and the presence of the other ancestral populations is very low or almost absent. Wild *Vitis vinifera* follows the classic phylogeographical pattern described in Hewitt, (2000) as “the richness of the south versus the purity of the north”. This pattern is explained by the demographic stability of the populations in southern refugia during the Quaternary glaciations, which led to greater polymorphism in the south and the loss of diversity experienced during the colonization of the north (Hewitt, 1996). In addition, the topography of the southern part of the peninsula favored allopatric differentiation between different populations, resulting in this well-known pattern of refugia within refugia, with greater diversity in southern areas (reviewed Gómez & Lunt, 2007).

Historically, the Iberian Peninsula has been studied as a refugium during the Pleistocene for different taxa, fish (Brito et al., 1997; Carmona et al., 2000; Machordom & Doadrio, 2001), invertebrates (Burban et al., 1999; Gómez et al., 2002), mammals (Pérez-Suárez et al., 1994; Branco et al., 2002), amphibians (Alexandrino et al., 2002), and reptiles (Paulo et al., 2001; Surget-Groba et al., 2001). Plants are no exception to this pattern, which has been investigated at both the genus and species levels. Some examples of this are: *Arabidopsis thaliana* (Picó et al., 2008), *Calluna vulgaris* (Rendell & Ennos, 2002), *Quercus* (Olalde et al., 2002; Petit et al., 2002; López de Heredia et al., 2007), *Quercus ilex* (Lumaret et al., 2002), *Hedera* (Grivet & Petit, 2002), *Pinus pinaster* (Salvador et al., 2000), *Pinus sylvestris* (Sinclair et al., 1999; Soranzo et al., 2000) and *Veronica aragonensis* (Padilla-García et al., 2021). According to Gómez & Lunt, (2007), the Iberian Peninsula, due to its high orographic complexity and strategic geographical position between the Atlantic and the Mediterranean, has a wide variety of microclimates throughout its distribution area, thus generating multiple and heterogeneous glacial refugia within the Iberian Peninsula.

Our study found high genetic diversity and rarity in wild grapevines within the Iberian Peninsula, mainly in the south and notably in the mountain ranges of Los Alcornocales, Sierra de Andújar, Sierra de Aracena, and Sierra de Hornachuelos. All these are within the Baetic mountain ranges, which have previously been proposed as a refugium for *Pinus pinaster* (Salvador et al., 2000), *Pinus sylvestris* (Sinclair et al., 1999; Soranzo et al., 2000) and *Quercus* (Petit et al., 2002) (reviewed in (Gómez & Lunt, 2007). Our results add to this cumulative knowledge, confirming that the south of the Iberian Peninsula has served as a refugium for plants during the Quaternary. This also supports the early hypotheses of (Levadoux, 1956) and re-stated by Grassi et al., (2008), who mentioned that during the Quaternary glaciations, wild grape populations were likely only found in refugia in the south of the Iberian Peninsula, Italy, and the Caucasus.

Regarding the demographic history and gene flow of wild grapevines within the Iberian Peninsula, our results suggest that the ancestral NA (K3) population derives from Middle Eastern individuals (notably from Georgia and Israel), which may have reached Iberia via a southern route through North Africa. East–west movements of both wild and cultivated grapevines have been previously reported (e.g. Freitas et al., 2021; Dong et al., 2023). However, despite early hypotheses proposing contacts between African and European grapevine populations, a clear Africa–Europe connection for wild grapevines has remained largely undetected in genomic studies. Notably, Arroyo-García et al., (2006), based on nine plastid loci, recovered African *V. sylvestris* as sister to Western European *V. sylvestris* (including Central European and Iberian populations), and identified the Iberian chlorotype A as the dominant chlorotype in African wild grapevines. Similarly, Laucou et al., (2018) found a subset of African and Iberian cultivated grapevines clustering within the same genetic group. Together, these findings support the hypothesis that the Strait of Gibraltar and adjacent regions may have functioned as a biogeographic corridor between North Africa and Europe at different times (Jaramillo-Correa et al., 2010; Husemann et al., 2014), a role previously documented in other plant lineages such as Linaria (Fernández-Mazuecos & Vargas, 2011; Blanco-Pastor & Vargas, 2013) and Cistus (Guzmán & Vargas, 2009; Fernández-Mazuecos & Vargas, 2010).

In this study, we evaluated demographic models to explain how and when this genetic contact occurred. Our results suggest a migration of grapevines from Africa to the Iberian Peninsula, with a continuous gene flow occurring at least since the Last Glacial Maximum (∼21,000 years ago) until the early Holocene (∼8,300 years ago), rather than gene flow pulses at specific time periods. Our findings support the ecological niche models presented by Dong et al., (2023), which indicate a niche differentiation between eastern and western wild grapevines during the Last Glacial Maximum (∼21,000 years ago), as well as the existence of a contact zone between both groups in the southern Iberian Peninsula. This evidence and our results support a natural, non-anthropogenic gene flow between Iberian and North African genotypes. Furthermore, other factors may have contributed to gene flow among wild grapevine populations. For instance, Debussche & Isenmann (1994) reported birds as seed dispersers of *Vitis* species in the Mediterranean region, while Terral et al., (2010) noted that the fruit ripening period of grapevines coincides with the migration of bird families such as Turdidae, facilitating seed dispersal through avian activity.

## Conclusion

The results of this study allowed us to identify the southern Iberian Peninsula as a hotspot of genetic diversity for the wild grapevines, as three ancestral populations or genetic groups were detected across this region. Likewise, it is important to highlight the “refugia within refugia” role of the Baetic mountain ranges within the Iberian Peninsula, given that four geographic contact zones among ancestral populations were identified in this area (Los Alcornocales, Sierra de Andújar, Sierra de Aracena, and Sierra de Hornachuelos), which can be proposed as micro refugia.

The genetic exchange detected between genotypes closely related to Middle Eastern lineages and likely migrating through Northern Africa and those originating in the Iberian Peninsula, was likely facilitated by climatic conditions from the Pleistocene (∼21,000 years ago) to the early Holocene (∼8,300 years ago) and has contributed additional genetic variability of wild grapevines in the region. This pattern is more consistent with natural, climate-driven connectivity than anthropogenic activities, suggesting that a substantial fraction of the observed genetic diversity predates the development of agriculture and grapevine domestication.

Finally, this work will serve as the basis for increasing the collection of Iberian wild vines. To this end, it will be necessary to continue identifying wild populations nearby high cluster diversity, and high heterozygosity and rarity, and to prioritize them for in situ protection and ex situ conservation before they are further eroded or lost (Lara et al., 2017; Iriarte-Chiapusso et al., 2017). This will enable the development of strategies and actions aimed at managing and promoting the conservation of the populations, taking into consideration its importance both as a component of biological diversity and as a genetic resource for cultivated grapevines in the current context of global change.

## Acknowledgements

We thank the Information Systems Unit of the Information Systems Area at the University of Cádiz for providing computational resources and technical support.

## Author Contributions

J.L.B.P. designed the study. J.J.R.F and J.J.S.B. data curation. J.J.R.F. analyzed the data. J.J.R.F. wrote the original draft. J.L.B.P. and J.J.S.B. review and edit the manuscript.

## Conflicts of Interest

The authors declare no conflicts of interest.

## Data Availability Statement

Genetic sequences were obtained from Dong et al., (2023), from BioProject PRJCA009314, file CRA006917, https://ngdc.cncb.ac.cn/gsa/browse/CRA006917. The dataset supporting this study is available in Dryad at https://doi.org/10.5061/dryad.sj3tx96k6.

Reviewers may access the data during peer review at: http://datadryad.org/share/LINK_NOT_FOR_PUBLICATION/4LbV3xd48jerp1FIkXhnYhw8PYmrjpdWkXuasu17vxM

## Notes

### Competing Interest Statement

The authors have declared no competing interest.

### Funding

- Junta de Andalucia: EMC21_00207

