## Appendix for "Southern Iberia as a hotspot of wild grapevine genetic diversity"

**Supporting Information**

**Appendix S1. Supplementary Tables**

**Table S1.** Sequences from the China National Center for Bioinformation, used for phylogenetic, genetic structure, and demographic history analyses. Obtained from Dong et al., (2023)

| Accession | ID | Locality |
|---|---|---|
| CRR499570 | PO161 | Portugal |
| CRR499572 | PO162 | Portugal |
| CRR499573 | PO163 | Portugal |
| CRR499575 | PO164 | Portugal |
| CRR499578 | PO165 | Portugal |
| CRR499580 | PO166 | Portugal |
| CRR499582 | PO167 | Portugal |
| CRR499584 | PO168 | Portugal |
| CRR499586 | PO169 | Portugal |
| CRR499588 | PO170 | Portugal |
| CRR499590 | PO171 | Portugal |
| CRR499592 | PO172 | Portugal |
| CRR499593 | PO173 | Portugal |
| CRR499595 | PO174 | Portugal |
| CRR499597 | PO175 | Portugal |
| CRR499599 | PO176 | Portugal |
| CRR499601 | PO177 | Portugal |
| CRR499603 | PO178 | Portugal |
| CRR499605 | PO179 | Portugal |
| CRR499607 | PO180 | Portugal |

| | | |
|---|---|---|
| CRR499609 | PO181 | Portugal |
| CRR499611 | PO182 | Portugal |
| CRR499613 | PO183 | Portugal |
| CRR499615 | PO184 | Portugal |
| CRR499617 | PO185 | Portugal |
| CRR498555 | 537 | Spain |
| CRR498461 | 538 | Spain |
| CRR498472 | 539 | Spain |
| CRR498475 | 540 | Spain |
| CRR498477 | 541 | Spain |
| CRR498483 | 542 | Spain |
| CRR498485 | 543 | Spain |
| CRR498487 | 544 | Spain |
| CRR498489 | 545 | Spain |
| CRR498491 | 546 | Spain |
| CRR498493 | 547 | Spain |
| CRR498495 | 548 | Spain |
| CRR498497 | 549 | Spain |
| CRR498499 | 550 | Spain |
| CRR498501 | 551 | Spain |
| CRR498503 | 552 | Spain |
| CRR498505 | 553 | Spain |
| CRR498507 | 554 | Spain |
| CRR498508 | 555 | Spain |

| | | |
|---|---|---|
| CRR498510 | 556 | Spain |
| CRR498512 | 557 | Spain |
| CRR498515 | 558 | Spain |
| CRR498517 | 559 | Spain |
| CRR498519 | 560 | Spain |
| CRR498521 | 561 | Spain |
| CRR498523 | 562 | Spain |
| CRR493851 | Spa32 | Spain |
| CRR493944 | Spa33 | Spain |
| CRR493946 | Spa34 | Spain |
| CRR493948 | Spa35 | Spain |
| CRR494314 | Spa36 | Spain |
| CRR494317 | Spa37 | Spain |
| CRR494319 | Spa38 | Spain |
| CRR494320 | Spa39 | Spain |
| CRR494322 | Spa40 | Spain |
| CRR494324 | Spa41 | Spain |
| CRR494326 | Spa42 | Spain |
| CRR494328 | Spa43 | Spain |
| CRR494330 | Spa44 | Spain |
| CRR494332 | Spa45 | Spain |
| CRR494334 | Spa46 | Spain |
| CRR494335 | Spa47 | Spain |
| CRR494337 | Spa48 | Spain |

| | | |
|---|---|---|
| CRR494339 | Spa49 | Spain |
| CRR494463 | Spa50 | Spain |
| CRR494848 | Spa51 | Spain |
| CRR494850 | Spa52 | Spain |
| CRR494852 | Spa53 | Spain |
| CRR494856 | Spa54 | Spain |
| CRR494858 | Spa55 | Spain |
| CRR494860 | Spa56 | Spain |
| CRR494863 | Spa57 | Spain |
| CRR494865 | Spa58 | Spain |
| CRR494867 | Spa59 | Spain |
| CRR494937 | Spa60 | Spain |
| CRR495519 | Spa61 | Spain |
| CRR495652 | Spa62 | Spain |
| CRR495654 | Spa63 | Spain |
| CRR495656 | Spa64 | Spain |
| CRR495658 | Spa65 | Spain |
| CRR495662 | Spa66 | Spain |
| CRR495664 | Spa67 | Spain |
| CRR495864 | Spa68 | Spain |
| CRR495865 | Spa69 | Spain |
| CRR496480 | Spa70 | Spain |
| CRR496482 | Spa71 | Spain |
| CRR496559 | Spa72 | Spain |

| | | |
|---|---|---|
| CRR496561 | Spa73 | Spain |
| CRR496563 | Spa74 | Spain |
| CRR496565 | Spa75 | Spain |
| CRR497174 | Spa76 | Spain |
| CRR497176 | Spa77 | Spain |
| CRR497859 | Spa78 | Spain |
| CRR497861 | Spa79 | Spain |
| CRR498463 | Spa80 | Spain |
| CRR498465 | Spa81 | Spain |
| CRR498466 | Spa82 | Spain |
| CRR498468 | Spa83 | Spain |
| CRR498471 | Spa84 | Spain |
| CRR498479 | Spa85 | Spain |
| CRR498481 | Spa86 | Spain |
| CRR498684 | Spa87 | Spain |
| CRR498686 | Spa88 | Spain |
| CRR498688 | Spa89 | Spain |
| CRR498690 | Spa90 | Spain |
| CRR498692 | Spa91 | Spain |
| CRR498855 | Spa92 | Spain |
| CRR498857 | Spa93 | Spain |
| CRR498859 | Spa94 | Spain |
| CRR499526 | Spa95 | Spain |
| CRR499527 | Spa96 | Spain |

| | | |
|---|---|---|
| CRR499529 | Spa97 | Spain |
| CRR493565 | IS97 | Israel |
| CRR493571 | IS103 | Israel |
| CRR493601 | IS135 | Israel |
| CRR493656 | IS191 | Israel |
| CRR495674 | IS95 | Israel |
| CRR496074 | GM34 | Germany |
| CRR496081 | GM38 | Germany |
| CRR499625 | GM15 | Germany |
| CRR497184 | GA3 | Georgia |
| CRR497251 | GA1 | Georgia |
| CRR497253 | GA4 | Georgia |
| CRR498784 | GA20 | Georgia |
| CRR499075 | GA10 | Georgia |
| CRR497515 | HU28 | Hungría |
| CRR497661 | AU84 | Austria |
| CRR498930 | Fre305 | Francia |
| CRR498543 | 576 | Francia |
| CRR498539 | 578 | Francia |
| CRR498535 | 579 | Francia |
| CRR498533 | 581 | Francia |

**Appendix S2. Supplementary Figures**

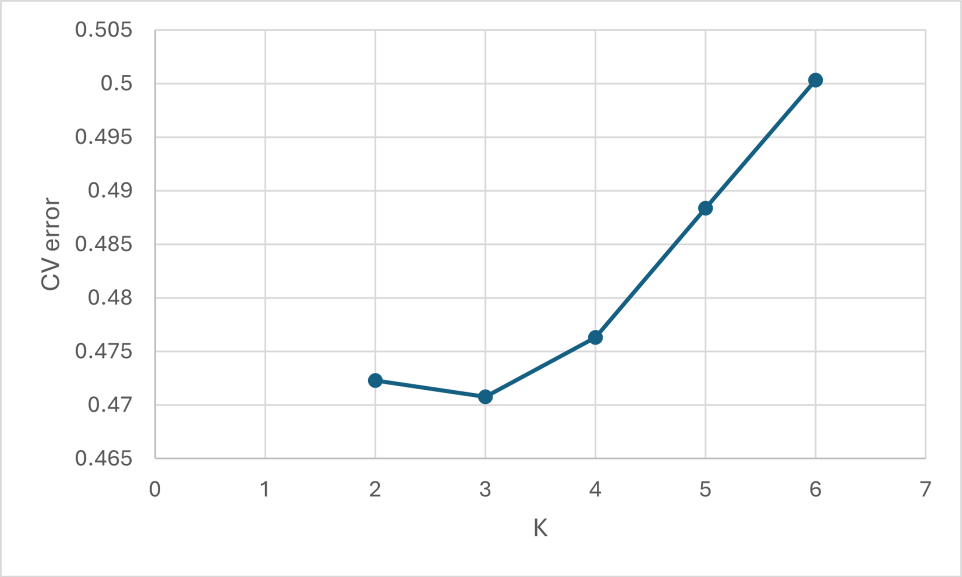


**Figure S1**. Cross-validation (CV) error graph for admixture analysis.

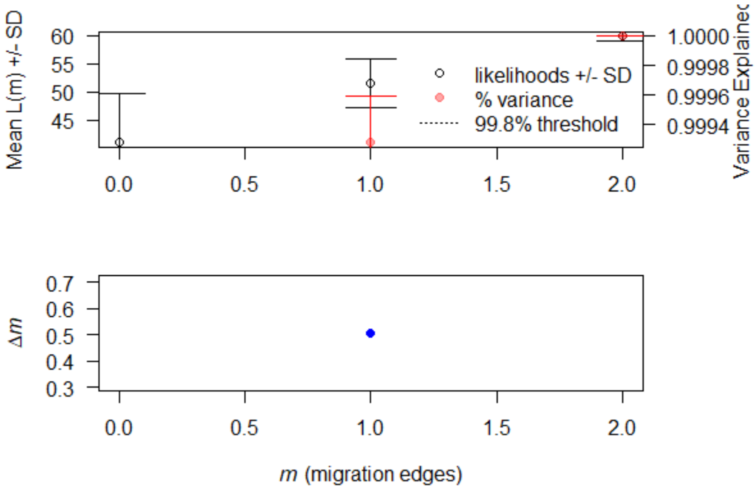


**Figure S2.** Selection of the optimal number of migration edges (m) using OptM. The upper panel shows the mean likelihood (black circles with standard deviation) and the proportion of variance explained (red dots) for models with 0 - 2 migration edges. Although the explained variance increases with m = 2, the main gain in model fit occurs when moving from m = 0 to m = 1. The lower panel represents the relative change in model fit (Δm), confirming that the largest increase occurs with the first migration edge. The optimal number of edges is considered to be m = 1, even though the absolute explained variance remains low.

**Data set**

http://datadryad.org/share/LINK_NOT_FOR_PUBLICATION/4LbV3xd48jerp1FIkXhnYhw8PYmrjpdWkXuasu17vxM
